# SMEW: An interactive multi-scale toolkit for cross-condition and network-based analysis of spatial metabolomics data

**DOI:** 10.64898/2026.04.27.721059

**Authors:** Eleanor C Williams, Heather Hulme, Aleksandr Zakirov, Daria Buszta, Gregory Hamm, Lucy Flint, Lovisa Franzén, Martina Olsson Lindvall, Marianna Stamou, Patrik Andersson, Jennifer Tan, Stephanie Ling, Irina Mohorianu

## Abstract

Spatial metabolomics, measured through mass spectrometry imaging (MSI), provides high-throughput, spatially resolved information on metabolite distributions within tissues, including endogenous metabolites and exogenous compounds. This offers a direct readout of cellular biochemical activity and phenotypes, not fully captured by transcriptomics or proteomic profiling. However, inferring biologically meaningful patterns from noisy, high-dimensional MSI data, particularly across multiple samples and complex experimental designs, remains challenging, and often requires substantial programming expertise.

Here we introduce SMEW (**S**patial **M**etabolomics **E**nhanced **W**orkflow), a flexible, interactive and shareable Shiny-based platform designed to enable code-free downstream analysis of spatial metabolomics MSI data. SMEW provides a unified environment for hierarchical analysis across bulk-, region- and pixel-level resolutions, allowing comparisons between experimental conditions like disease or treatment groups while highlighting coherent metabolic patterns and linking these patterns to biological pathways. The workflow leverages local spatial covariation to robustly summarise MSI data through dimensionality reduction, clustering and identification of spatially variable metabolites. In addition, metabolite co-localisation and covariation network analysis, together with spatially resolved pathway enrichment facilitate the biological interpretation of cross-condition datasets within a single integrated interface.

SMEW is applicable across MSI technologies and mass resolutions, as illustrated through case studies on DESI and MALDI-ToF datasets from lung, liver, and kidney. By complementing existing MSI processing and visualisation tools with an accessible, multi-sample, and biologically interpretable analysis framework, SMEW enables functional, flexible, rigorous and intuitive exploration of spatial metabolomics datasets.

**Graphical Abstract:** 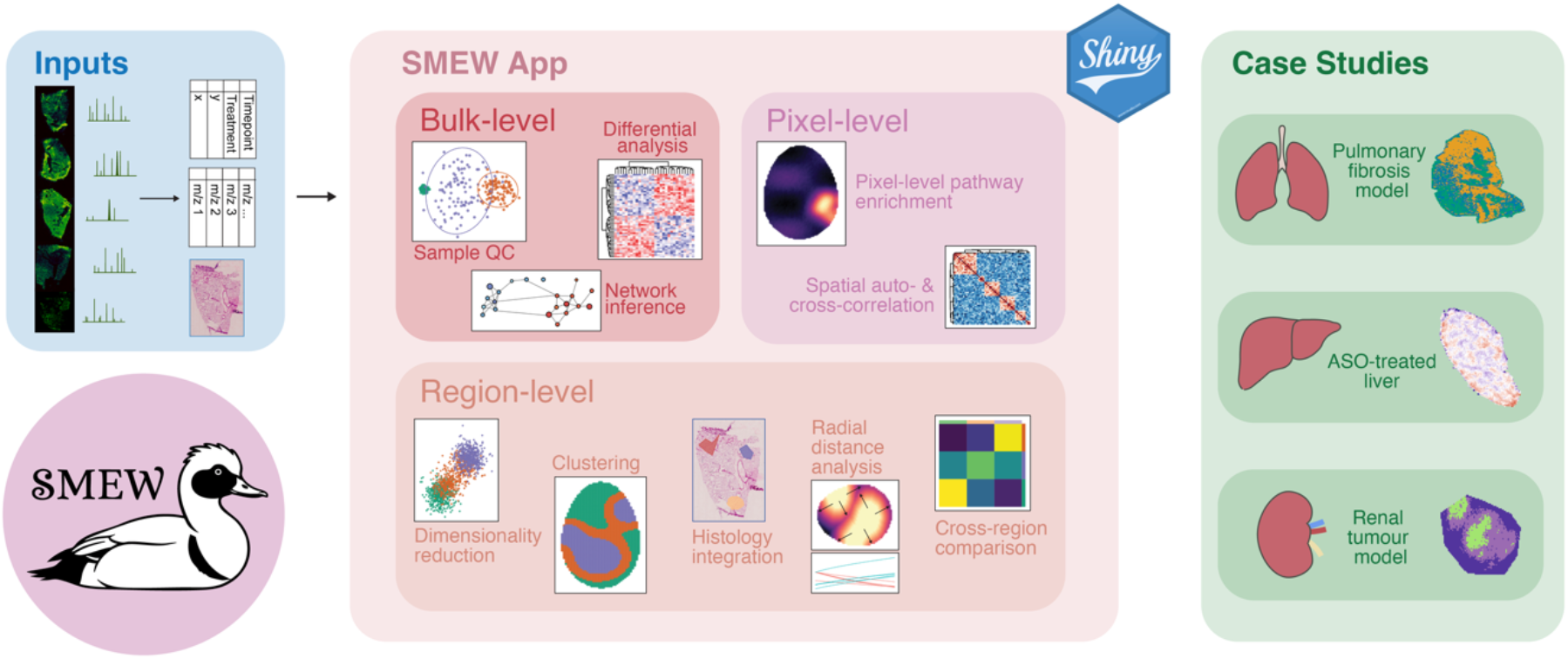

**Key Points:**

- SMEW provides a flexible, interactive and shareable Shiny-based platform designed to enable code-free downstream analysis of spatial metabolomics MSI data
- The SMEW framework enables hierarchical analysis at bulk-, region- and pixel levels within a unified framework without relying on extensive programming expertise
- The pipeline integrates spatially aware clustering, pathway analysis and identification of metabolite co-localisation modules
- The workflow facilitates flexible comparison of multi-sample experimental conditions through multivariate modelling, differential testing and covariation networks to study treatment- and disease-associated metabolite dynamics
- SMEW has been applied to interrogate diverse biological questions, including characterising disease-associated remodelling in a mouse bleomycin model of pulmonary fibrosis, exploring the therapeutic index of antisense oligonucleotides in the liver and assessing metabolic heterogeneity in a small molecule-treated mouse renal tumour model

## Introduction

Rapid advances in single-cell and spatial multi-omics technologies have transformed our ability to infer mechanistic and regulatory processes within tissues. Although historically work in this area has been driven by transcriptomic profiling, there is increasing interest in capturing the biochemical state of biological systems by measuring small molecules via metabolomics. Mass spectrometry imaging (MSI) has emerged as a powerful approach for studying the spatial localisation of metabolites, lipids and pharmacological compounds in cells, organs and histologically defined regions of interest within tissues^1,2^. MSI provides a snapshot of metabolite distributions at the time of tissue collection; while this reflects steady-state abundances rather than directly measuring metabolic flux, inferring spatial metabolite distribution patterns can still reflect dynamic metabolic processes and provide information on metabolic reprogramming due to disease and injury^3,4^. This is achieved by dividing tissues into grids of pixels and acquiring a mass spectrum per pixel. Different ionisation methods, such as matrix-assisted laser desorption/ionisation (MALDI), desorption electrospray ionisation (DESI) and secondary ion mass spectrometry (SIMS), can be used, and these are coupled with various mass analysers depending on desired analyte mass ranges, spatial resolutions, mass accuracies and time requirements^1,2^. Spatial metabolomics through MSI has the potential to offer key insights across biological domains, e.g. to probe tumour heterogeneity^5^, treatment and exposure response^6^, or to uncover the spatial metabolic organisation of organs^7^.

In recent years, several tools have been developed to enable the exploration of spatial metabolomics data, addressing different aspects of the MSI workflow. rMSI^8,9^ and MALDIquant^10^ focus on early-stage processing of data in standard imzML format, including peak selection and normalisation, but offer limited support for integrated downstream statistical analyses. Interactive environments such as the commercial SCiLS™ Lab software by Bruker and non-commercial tools including MSI-Explorer^11^ and the Galaxy-MSI^12^ platform, have improved accessibility of MSI analysis through graphical interfaces. These platforms provide insightful visualisation and segmentation capabilities, however integration of multi-sample statistical modelling, pathway enrichment and network-based biological interpretation across complex experimental designs often requires additional tools. More detailed downstream analysis is offered by the Cardinal package, a comprehensive R-based framework for spatially aware statistical modelling and segmentation, including principal component analysis, spatially informed clustering and supervised classification approaches tailored to MSI data. ShinyCardinal^13^ enables a subset of these analyses to be performed interactively in a code-free setting. While these frameworks offer a range of analysis options, translating MSI data into higher-level biological insight and bringing together pseudobulk- (i.e. aggregated to sample-level data), region- and pixel-level analyses often require multiple disconnected workflows. Consequently, there remains an unmet need for a unified, user-friendly analytical framework that complements existing MSI toolkits by enabling multi-scale and cross-sample statistical modelling, spatially informed analyses and network-level interpretation within a single, accessible interface.

To address these challenges, we developed SMEW (**S**patial **M**etabolomics **E**nhanced **W**orkflow), an interactive Shiny application designed to complement existing MSI processing and modelling tools while facilitating downstream, code-free analysis of spatial metabolomics data. The pipeline is open source and available on GitHub with extensive documentation and comprehensive case studies. SMEW focuses on orchestrating and integrating spatially informed analyses across multiple samples and experimental covariates, enabling users to move beyond single-sample exploration toward dataset-wide inference. The platform supports hierarchical analysis across bulk-level comparisons, regional cluster analysis, and pixel-level inference within a consistent conceptual pipeline. It also allows users to incorporate intermediate results generated by other platforms. SMEW further extends spatial metabolomics analysis toward biological interpretation through features such as pathway-level summarisation, spatially resolved pathway enrichment and covariation network analysis to probe regulatory relationships^14^. Multi-omics integration, e.g. with transcriptomic or proteomic data, is also possible within SMEW, offering advantages over existing tools. Leveraging methodological concepts from spatial transcriptomics, SMEW enables real-time quality control, visualisation of spatial peak distributions and exploration of metabolite co-localisation across samples and experimental groups in a code-free, robust, scalable and interactive environment.

## Results

SMEW (**S**patial **M**etabolomics **E**nhanced **W**orkflow) provides an interactive environment that integrates dataset-wide, spatially informed analyses into a single coherent workflow, enabling users to move seamlessly from global comparisons to localised spatial inference without requiring programming expertise. The framework is organised around three tightly linked analytical components performing pseudobulk-level, region-level and pixel-level analysis, mirroring complementary biological scales of interpretation (**Fig. 1**).

**Figure 1.**
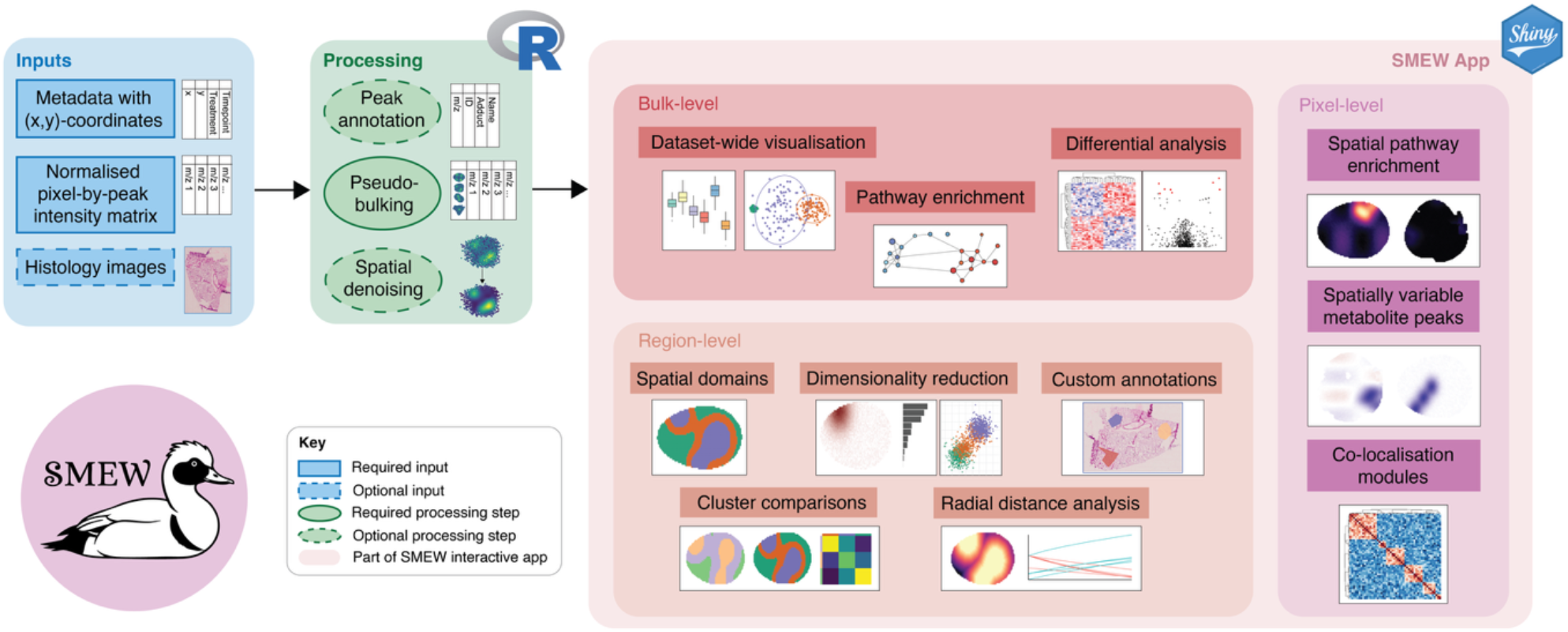
SMEW Workflow: Schematic representation of the SMEW pipeline, illustrating the hierarchical and modular analysis of mass spectrometry imaging (MSI) data. MSI datasets from different platforms can be imported and pre-processed, then SMEW creates an interactive app to enable flexible exploratory analyses at the bulk, region, and pixel levels.

Here we provide a detailed overview of the full SMEW pipeline, illustrating the flexibility and versatility of the workflow across multiple biological scales and experimental designs. We first illustrate the analysis steps using a dataset from a mouse model of pulmonary in negative ion mode, with samples collected at 7 and 21 days post-bleomycin treatment to model early inflammation in the lung and established tissue damage and fibrosis, respectively^15,16^. We then showcase SMEW’s versatility by applying the workflow to a DESI dataset investigating hepatotoxicity of antisense oligonucleotides and a MALDI-ToF dataset of renal tumour response to treatment^6^. Together, these examples highlight SMEW’s ability to integrate multi-sample analyses, capture spatially resolved metabolic patterns and accommodate diverse MSI technologies and biological contexts.

### Overview of SMEW interactive pipeline illustrated on a bleomycin-treated mouse lung case study

#### Inputs and pre-processing

The SMEW pipeline requires a peak-by-pixel format as input, which can be generated using state-of-the-art tools such as the commercial tool SCiLS™ and open-source methods MZmine^17^, MALDIquant^10^ or Cardinal^18^, all of which accept the open-source imzML format as input. Input data should be pre-normalised (e.g. using Total Ion Count or Root Mean Square), ensuring that intensities are comparable across pixels. SMEW provides an additional, optional denoising and spatial smoothing step, which combines signals from neighbouring pixels with similar intensity patterns using principal component-based correlation. Alternatively, externally smoothed data can be incorporated^18,19^.

Putative peak annotation can be performed directly within SMEW by comparing measured *m/z* values with expected metabolite masses within a user-specified ppm threshold using a user-selected database such as KEGG^20^, HMDB^21^ or LipidMaps^22^, or by integrating external tools such as Cardinal^18^, rMSIannotation^23^ and METASPACE^24^ which use additional information such as isotopic patterns, fragmentation and spatial context to refine annotations. The intensity matrix, metadata and peak annotations are then used to generate the SMEW app. To support an initial assessment of the data, the app begins with introductory tabs enabling interactive exploration of the annotation table and searching for metabolites of interest, as well as visualisation of spatial gradients of individual peaks, combinations of peaks and user-supplied pixel-level metadata. For example, in a model of bleomycin induced lung fibrosis, peak *m/z 303*.*23295* (FA 20:4 Arachidonic acid) shows clear spatial heterogeneity and treatment-related changes across samples, highlighting SMEW’s ability to capture treatment-specific spatial patterns.

#### Pseudobulk-level analysis highlights condition-specific metabolite changes

To explore global spatially agnostic trends and further assess dataset quality across samples, SMEW’s pseudobulk-level tabs provide quality control summaries and enable statistical comparisons of MSI data across experimental conditions. These include principal component analysis (PCA) and partial least squares discriminant analysis (PLS-DA) to evaluate replicate consistency, detect batch or slide effects and group separation, alongside visualisation of individual peak intensities across samples. In the bleomycin-treated mouse lung dataset^15,16^ (*n*=24), PCA reveals some separation of treatment groups, further emphasised by PLS-DA (**Fig. 2a**). SMEW enables the identification of metabolite peaks inducing this separation; treatment groups differences are driven by peaks such as *m/z 121*.*02969* (Benzoate), *m/z 129*.*01922* (Itaconate) and *m/z 465*.*30331* (Cholesterol sulphate) in the positive PLS-DA Component 1 direction, towards the control samples, and *m/z 365*.*34273* (FA 24:1 Tetracosenoic acid) and *m/z 337*.*31232* (FA 22:1 Erucic acid) in the negative direction towards the bleomycin samples, reflecting lipid remodelling. Further inspection of the intensities of top peaks using box plots (**Fig. 2b**) reveals a gradient of intensity from the two control groups through to bleomycin treatment after 7 days and finally after 21 days.

**Figure 2.**
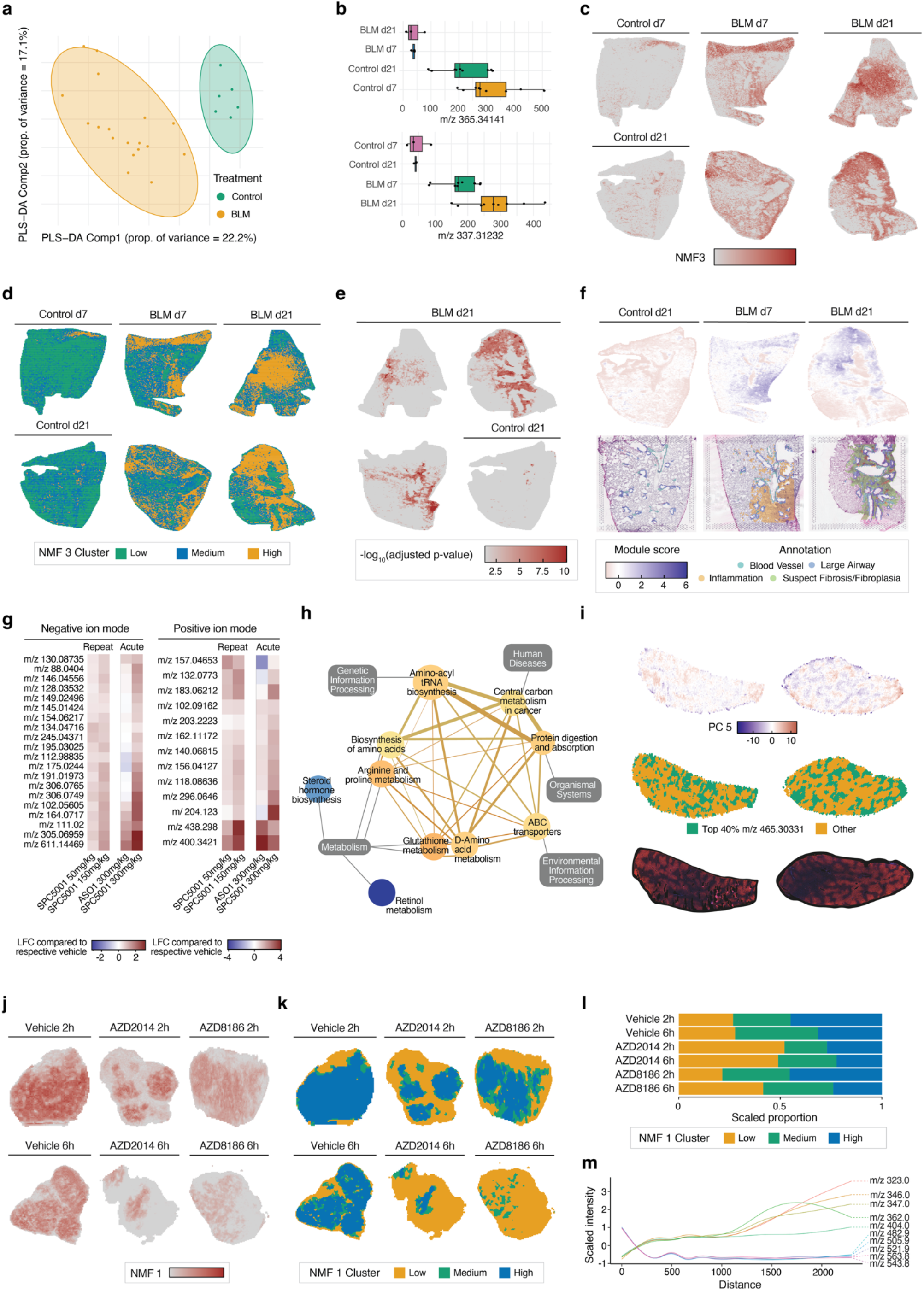
Application of SMEW across three spatial metabolomics case studies. (a-f) Bleomycin-induced lung fibrosis (DESI MSI). (a) Bulk-level principal component analysis (PCA) summarising global metabolic variation across bleomycin-treated (BLM) and control samples. (b) Boxplots showing the distribution of intensities for selected top PLS-DA metabolite peaks at the bulk level across BLM and control treatment groups 7 days and 21 days post-exposure. (c) Spatial view of non-negative matrix factorisation (NMF) factor 3, highlighting regional metabolic heterogeneity specific to BLM samples. (d) Spatial view of clusters derived from thresholding on NMF factor 3, categorised into low, medium and high intensity regions. (e) Pixel-level pathway analysis of the alpha-linolenic acid metabolism pathway visualised as −log_10_(BH-adjusted p-value) across the tissue section. (f) Spatially variable metabolite co-localisation module scores (aggregated by mean across scaled module peaks) displayed alongside pathology annotations on adjacent tissue sections overlaid on corresponding H&E staining. **(g–i) ASO-induced liver toxicity (DESI MSI)**. (g) Heatmaps showing log_2_ fold change (LFC) between treated and respective controls (acute or repeat study) for top differentially abundant metabolite peaks comparing SPC5001 and control samples in negative (left) and positive (right) ion modes. (h) Pathway network showing over-representation analysis on differentially abundant metabolite peaks comparing SPC5001 and control samples. Node colour represents the increase (red) or decrease (blue) in activity of each pathway, and edges represent overlapping metabolites between pathways. Pathways are also labelled by KEGG categories in grey rectangles. (i) Liver zonation analysis showing spatial distribution of PC5, top 40% cholesterol sulphate (*m/z 465*.*30331*) pixels and Cyp2e1 staining on adjacent sections, highlighting periportal and pericentral metabolic differences. **(j–m) Small molecule-treated renal tumours (MALDI-ToF MSI)**. (j) Spatial view of NMF Factor 1 in selected samples, illustrating spatial and treatment tumour heterogeneity. (k) Spatial view of NMF Factor 1-based regions in selected samples, where “Low” corresponds to the bottom 40%,”High” the top 40% and “Medium” the middle 20% by NMF Factor 1. (l) Proportion of NMF Factor 1-based regions within each treatment group, scaled by region size to reflect relative enrichment of metabolic states. (m) Radial distance line plots showing scaled intensities of metabolite peaks most positively and negatively correlated with distance from the NMF Factor 1 high region in AZD2014-treated tumours.

SMEW further enables the identification of significantly changing peaks across conditions using t-tests, Wilcox rank sum tests, ANOVA, or Kruskal-Wallis with post-hoc comparisons and user-selected significance thresholds. The results can then be explored through interactive tables, visualised using volcano plots and summarised in heatmaps. Pathway over-representation analysis further contextualises metabolite changes, with results displayed through tables, hierarchical pathway grouping and network diagrams indicating shared metabolites and direction of change. Covariation network analysis at the pseudobulk level facilitates the identification of coordinated metabolic modules. SMEW further supports the incorporation of multi-omics data e.g. from transcriptomics or proteomics, to generate integrated networks, highlighting the pipeline’s unique capacity to extend single-peak insights into system-level patterns.

In our mouse lung case study^15,16^, SMEW’s differential intensity analysis and heatmap summaries enable us to observe increased levels of several prostaglandins (*m/z 405*.*20581, 387*.*19378, 333*.*2071*) and FA 20:4 Arachidonic acid (*m/z 303*.*23295*) in bleomycin-treated samples, a hallmark of inflammatory responses, consistent with previous findings in bleomycin-induced pulmonary fibrosis^25,26^. We further observe that Glutathione (*m/z 306*.*07653*) increases 7 days post-bleomycin challenge during the acute inflammatory phase following injury then reduces at 21 days, while FA 24:1 Tetracosanoic acid (*m/z 365*.*34273*) and FA 22:2 Docosadienoic acid (*m/z 335*.*29683*) increase slightly at 7 days then further increase at 21 days as fibrosis becomes evident. SMEW’s multi-comparison tab allows us to further probe changes across the time course using ANOVA, highlighting FA 4:0;O Hydroxybutanoic acid (*m/z 103*.*0401*) and FA 20:1 Icosenoic acid (*m/z 309*.*28026*) as specific to day 7’s early metabolic stress and day 21’s later lipid remodelling post-bleomycin treatment respectively. Pathway analysis highlights an increase in arachidonic acid metabolism, linoleic acid metabolism and biosynthesis of unsaturated fatty acids; this highlights the demand for new membrane lipids due to the tissue injury, inflammation and fibrotic remodelling that may be caused by the treatment. Covariation networks comparing samples 7 days and 21 days post-bleomycin exposure further reveal conserved lipid modules centred on FA 20:4 Arachidonic acid (*m/z 303*.*23295*) and FA 18:2 Linoleic acid (*m/z 279*.*23186*), with consistent connections to related polyunsaturated fatty acids such as FA 20:3 Icosatrienoic acid (*m/z 305*.*24751*). In addition, stage-specific covariation patterns were observed, demonstrating SMEW’s ability to capture dynamic changes in metabolite networks across post-exposure phases.

#### Region-level analysis identifies data-driven spatial domains of interest

While pseudobulk analyses capture global patterns across samples, SMEW also enables more granular, spatially-resolved analyses of metabolite distributions through its region-level capabilities. SMEW supports multiple approaches to define spatially-resolved regions of interest, including dimensionality reductions such as non-negative matrix factorisation (NMF), PCA, optionally layered with UMAP to find key axes of variation and the peaks driving them (**Fig. 2c**), as well as clustering methods (**Fig. 2d**) such as k-means, Louvain community detection and spatially-informed BayesSpace^27^. Cluster identification can incorporate batch correction via Harmony^28^ to account for and minimise technical differences between samples, such as between slides or lab environments. Spatial clusters identified within individual samples can be combined across the dataset through hierarchical clustering to enable cross-condition analysis.

Regions of interest can also be defined using thresholding on individual peak intensities or combinations of peaks, or through interactive selection, including co-registration with histological images (e.g. H&E staining) provided alongside SMEW. This allows histological annotations to be overlaid with metabolite information to assess their enrichment in areas of pathological interest.

Regions identified through the Region-level tabs, alongside any user-defined regions, can be spatially smoothed to improve cluster stability and interpretability, following strategies established in spatial transcriptomics^29^. Regions can then be quantitatively compared against each other (e.g. Jaccard similarity index and element centric similarity^30^), allowing users to assess where spatially defined metabolic regions overlap or differ, providing insight into the organisation of metabolic states within the tissue. Marker peaks can then be identified per cluster and optionally stratified by metadata. SMEW further supports region-level covariation network analysis using GENIE3^31^, enabling exploration of regulatory relationships across spatial domains and experimental conditions. Finally, radial distance analysis^32^ quantifies metabolite gradients relative to selected regions, enabling comparison of metabolite intensities in the immediate microenvironment versus more distal areas.

In the bleomycin lung example, PCA scores shown spatially revealed that key metabolic variation in the dataset is driven by treatment group, while NMF highlighted factors which were consistently high across all treated samples (e.g. NMF2) and specific to early inflammation (NMF1, day 7). NMF1 is driven by TCA cycle intermediates including Malate (*m/z 133*.*01425*) and Citrate (*m/z 191*.*01973*), which could suggest metabolic reprogramming in the immune landscape during the early inflammatory phase after bleomycin exposure. NMF3 shows more spatial heterogeneity (**Fig. 2c**) and was used to form clusters. By quantile-splitting NMF3 into High (top 25%), Medium (middle 50%) and Low (bottom 25%) pixel-level groups (**Fig. 2d**) and comparing against k-means clusters (k=4), we identified Cluster 1 as overlapping with High NMF3 regions and the histologically-assessed suspect fibrosis areas, while Cluster 2 overlapped with Low NMF3. Importantly, these regions are derived directly from spatial metabolite patterns, and this analysis does not require histopathological annotation, allowing metabolically distinct tissue regions to be identified in a data-driven manner, even when histology is unavailable. Radial distance analysis from Cluster 1 revealed localised enrichment of long-chain fatty acids and lipid oxidation products (*m/z 281*.*24751*, FA 18:1 Octadecenoic acid; *m/z 309*.*28026*, FA 20:1 Icosenoic acid; *m/z 171*.*10256*, FA 9:1;O 9-Oxononanoic acid), reflecting spatially restricted metabolic remodelling in fibrotic tissue. A voting scheme based on these peaks confirms their localisation to day 21 bleomycin-treated tissues in areas labelled as fibrotic. Together, these analyses illustrate how SMEW integrates complementary region-identification strategies within a single, interactive framework, enabling systematic comparison of spatial domains and their associated metabolic signatures across samples and conditions.

#### Pixel-level analysis reveals spatially localised metabolic programs and co-variation patterns

To fully exploit the spatial resolution of MSI data, SMEW offers pixel-level analysis approaches to identify localised metabolic changes across tissue sections, thus complementing pseudobulk- and region-level results. By explicitly modelling spatial neighbourhoods, these analyses capture heterogeneity that may be restricted to specific anatomical or pathological regions, such as focal disease lesions or areas of local drug accumulation.

SMEW implements a sliding window neighbourhood approach for spatial pathway enrichment, in which each pixel and its spatial neighbours are treated as local replicates and compared against pseudobulk control samples. Differentially abundant metabolites identified in these local comparisons are subsequently subjected to pathway over-representation analysis (ORA), enabling inference of spatially resolved pathway activity (**Fig. 2e**). Mapping pathway-level significance across the tissue reveals gradients of metabolic changes and assists in the interpretation of spatial domains, particularly in heterogeneous disease contexts. In the bleomycin mouse lung model, SMEW’s pixel-level enrichment highlighted several pathways with localised enrichment in treated samples, including alpha-Linoleic acid metabolism and Inflammatory mediator regulation of TRP channels, consistent with lipid remodelling and chronic inflammation in fibrosis.

Beyond pathway-level inference, SMEW also enables the identification of spatially variable metabolite peaks with distinct intensity patterns in space and modules of co-localised metabolites. Spatial autocorrelation is computed using a fast approach implemented based on the semla^32^ package for spatial transcriptomics analysis, quantifying the similarity between each pixel and its neighbours via a ‘spatial lag’ vector. Spatial autocorrelation metrics can be compared across samples and conditions using dot plots, upset plots and bar plots, incorporating bulk-level differential analysis information to highlight condition-specific spatial organisation. To further characterise coordinated spatial patterns, SMEW extends the spatial cross-correlation metric from MERINGUE^33^ to multi-sample MSI datasets to identify modules of metabolite peaks with similar spatial distributions or metabolite niches. These modules can be visualised using clustered heatmaps and co-localisation networks and projected back into tissue space to reveal coherent metabolic domains.

In the bleomycin dataset, spatial autocorrelation analysis identified a peak module with increased intensity in treated samples at both day 7 and 21, spatially aligned with regions of inflammation and suspect fibrosis (**Fig. 2f**). This modules includes several lipid species consistent with bleomycin-induced lipid remodelling, including FA 18:2 Linoleic acid (*m/z 279*.*23186*), FA 18:1 Octadecenoic acid (*m/z 281*.*24751*), FA 20:4 Arachidonic acid (*m/z 303*.*23295*), FA 22:6 Docosahexaenoic acid (*m/z 327*.*23186*), and oxidised derivatives like FA 18:3;O 13-OxoODE (*m/z 293*.*21263*), suggesting localised accumulation of fatty acids and lipid oxidation products, consistent with sustained oxidative stress at both days 7 and 21 post-challenge. While these metabolites are also detected in pseudobulk- and region-level analyses, SMEW’s pixel-level modules reveal where and how they co-localise, providing insights into spatially restricted metabolic changes and gradients of intensity that pseudobulk and discrete clustering cannot resolve.

Together, these pixel-level analyses illustrate how SMEW integrates pathway-level inference and spatial covariation analysis within a unified framework, enabling systematic identification of treatment-specific, spatially localised metabolic programmes and refining the characterisation of disease heterogeneity. We next present two additional case studies to underline SMEW’s generalisability across tissues, technologies and experimental designs.

### Applying SMEW to characterise ASO-induced liver injury

To demonstrate SMEW’s applicability and generalisability to different tissues, experimental designs and MSI technologies, we applied the workflow to a DESI MSI dataset investigating antisense oligonucleotide (ASO)-induced liver injury in mice. ASOs are short single-stranded deoxyribonucleotides (~18-30nt) designed to be complementary to their target in order to induce RNase H-dependent transcript degradation or alter splicing processes. However, ASOs have the potential to induce toxicity, particularly in the liver, where ASO uptake is elevated^34,35^. This dataset therefore provides a biologically relevant setting to showcase the SMEW framework’s ability to disentangle dose-, study- and region-specific metabolic effects.

The dataset (*n*=40) comprises liver sections from two independent studies. In the ‘Acute’ study (*n*=16), mice received single subcutaneous administrations of vehicle (saline) and two ASOs of similar chemistry and design at 300 mg/kg and were euthanised 3 days later. ASO1 is a liver safe ASO in this study, while SPC5001 shows medium liver toxicity^36^. In the ‘Repeat’ study (*n*=24), mice were treated subcutaneously once weekly for six weeks with vehicle (saline) or SPC5001 at 50 mg/kg or 150 mg/kg dose levels for 6 weeks. For DESI MSI acquisition, serial tissue sections from each sample were analysed separately in positive and negative ionisation modes, with samples from different treatment groups randomly distributed across three slides per polarity, introducing technical variation representative of real-world experimental designs.

Using SMEW’s pseudobulk-level analysis, we first assessed global variation across samples and studies per polarity. PCA revealed partial separation by treatment group alongside pronounced slide-associated effects. This highlighted a typical scenario where batch effects can influence clustering, motivating the use of the batch correction provided in the SMEW region-level analysis. Indeed, initial clustering of repeat-dose samples (positive detection ion mode) without batch correction produced clusters dominated by individual slides. By applying Harmony batch correction^28^ within SMEW’s region-level analysis module, technical variation across slides was significantly reduced, resulting in spatial clusters that were more evenly distributed across slides and strongly aligned with treatment groups. This highlights how SMEW enables systematic assessment of dataset quality and slide-specific effects, and especially its capacity to integrate multi-slide MSI datasets while preserving biologically meaningful structure.

SMEW was then used to perform bulk-level differential analysis between vehicle- and SPC5001-treated samples across studies and doses, identifying metabolite peaks that consistently increased under ASO exposure (**Fig. 2g**). Across both ion modes, the strongest changes were observed in the acute 300 mg/kg group, consistent with rapid, severe metabolic disturbance in acute toxicity. These included metabolites linked to oxidative stress (e.g., Glutathione disulphide, *m/z 611*.*14469*; Glutathione, *m/z 306*.*0765*) in negative ion mode and mitochondrial function (e.g., Acetylcarnitine, *m/z 204*.*123*) in positive ion mode. In repeat-dose samples, SMEW identified a subset of metabolites, including Creatine (*m/z 132*.*0773*) in positive ion mode, showing larger log fold changes than in the acute setting, while other features (e.g. long-chain Acylcarnitines (*m/z 400*.*3421, m/z 438*.*298*) in positive mode, amino acids (e.g., L-Glutamate, m/z 146.04556; L-Leucine, m/z 130.08735) in negative mode and TCA cycle intermediates like Citrate (*m/z 191*.*01973*) in negative ion mode exhibited consistent treatment-associated changes across both acute and repeat dose SPC5001 groups. These patterns illustrate how SMEW captures both dose-specific and shared metabolic responses across multiple studies.

To go a step further towards biological interpretation, SMEW’s pathway-level enrichment module highlighted coordinated shifts in metabolic programs under SPC5001 exposure (**Fig. 2h**). For example, treatment-associated changes in glutathione species in negative ion mode were reflected in enrichment of glutathione metabolism, while changes in amino acids aligned with pathways such as arginine and proline metabolism and broader amino acid biosynthesis. Enrichment of the aminoacyl-tRNA biosynthesis pathway was also detected, consistent with cellular stress responses associated with mitochondrial dysfunction. Together, these analyses demonstrate how SMEW bridges metabolite-level changes and pathway-level interpretation in a dataset-wide and condition-aware manner.

Beyond treatment-driven effects, SMEW also enabled the identification of anatomical organisation within the liver. Using pixel-level PCA (**Fig. 2i**) and marker-based segmentation on negative ion mode data (**Fig. 2i**), example tissues from one block across vehicle, ASO1 and SPC5001 groups were partitioned into periportal- and pericentral-like regions. These regions showed spatial patterns consistent with Cyp2e1 staining on adjacent sections (**Fig. 2i**), a marker associated with pericentral hepatocytes. Radial distance analysis on periportal-like regions revealed metabolite distributions consistent with known zonal metabolic profiles. Periportal areas were enriched for metabolite peaks linked to lipid transport and fatty acid metabolism including Cholesterol sulphate (*m/z 465*.*30331*), FA 14:0 Tetradecanoic acid (*m/z 227*.*20217*) and multiple polyunsaturated fatty acids such as FA 22:6 Docosahexaenoic acid (*m/z 327*.*23186*) and FA 18:3 Octadecatrienoic acid (*m/z 277*.*21621*), consistent with the known role of periportal hepatocytes in lipid uptake and oxidative metabolism. In contrast, pericentral regions showed higher relative abundance of metabolites linked to oxidative stress, lipid oxidation and inflammatory lipid signalling, such as leukotriene-like (*m/z 335*.*22185, m/z 337*.*23964*) and prostaglandin-like (*m/z 351*.*2177*) species, in line with the enrichment of xenobiotic metabolism and oxidative processes in pericentral hepatocytes. This SMEW-enabled segmentation approach provides a foundation for studying zonation-specific hepatic metabolic disruption, including the loss of zonation, such as in MASLD^37^.

Collectively, this case study demonstrates the efficiency of incremental analyses within SMEW and the added value of pseudobulk, region-level and pixel-level perspectives within a single interactive framework for dissecting complex MSI datasets. Moreover, by enabling batch-aware integration, multi-study comparisons, pathway-level interpretation and spatially resolved analysis, SMEW provides a flexible and generalisable approach for characterising drug-induced toxicity and tissue-specific metabolic organisation across diverse experimental settings.

### Explorative analysis of small molecule-treated renal tumours through SMEW

To further illustrate SMEW’s ability to interrogate complex experimental set-ups and its generalisability across MSI platforms, we applied the workflow to a negative ion mode MALDI-ToF MSI dataset of a mouse renal tumour model investigating early metabolic responses to targeted pathway inhibition (Ling *et al*. ^6^). This study (*n*=25) compared tumours treated *in vivo* with AZD2014, an mTORC1/2 inhibitor, AZD8186, a PI3Kβ inhibitor, or vehicle controls, assessed at 2 hours and 6 hours post-treatment. The experimental design thus captures both pathway-specific effects and early spatial heterogeneity in tumour response. This dataset also allows us to explore the types of analysis that can be performed in SMEW, even with low mass resolution data, where annotation to known metabolites and pathways can be challenging.

Using SMEW’s pixel-level dimensionality reduction, NMF revealed a factor (NMF1) exhibiting pronounced spatial heterogeneity within tumours (**Fig. 2j**) as well as systematic variation across treatment groups, reflecting SMEW’s ability to capture treatment-associated metabolic gradients directly at pixel resolution. NMF1 scores across all samples were subsequently used to define metabolic clusters (**Fig. 2k**), enabling a spatial stratification of tumour regions according to inferred metabolic activity. The resulting clusters showed the highest proportion of ‘High’ NMF1 pixels in Vehicle controls and AZD8186-treated tumours at 2 hours post-treatment, while AZD2014-treated tumours and tumours at 6 hours post-AZD8186 treatment exhibited the lowest NMF1 levels (**Fig. 2l**).

Further inspection of NMF1 loadings reveals a strong contribution from ATP (*m/z 505*.*9*) and ADP (m/z *426*.*0*), supported by the high similarity in spatial patterns between ATP intensity and NMF1 scores. The spatially localised reduction in ATP and ADP suggests decreased energy metabolism in affected tumour regions, with a more delayed effect observed for AZD8186. Radial distance analysis in AZD2014-treated tumours (**Fig. 2m**) also highlights that ATP intensity inversely correlates with distance from NMF1-high areas, demonstrating how complementary SMEW analysis modules can help to build up a picture of the metabolomic landscape.

Together, this case study highlights SMEW’s flexibility and robustness across MSI platforms and biological contexts. SMEW seamlessly accommodates both high-resolution DESI datasets and lower-resolution MALDI-ToF data, enabling coherent analysis despite differences in mass accuracy and feature complexity. By integrating pixel-level dimensionality reduction, spatial clustering, and radial distance analysis, SMEW reveals metabolically distinct tumour subregions and microenvironments corresponding to differential therapeutic response, consistent with previously published data^6^. This ability to resolve spatially confined metabolic adaptations provides a powerful foundation for interpreting heterogeneous drug responses and for guiding further mechanistic and translational investigations in oncology.

## Conclusion

Here, we introduced SMEW, an interactive platform for flexible and tailored analysis of spatial metabolomics MSI data. Through three case studies spanning DESI-Orbitrap datasets (a bleomycin model for idiopathic pulmonary fibrosis and a study of ASO-induced liver toxicity) and a MALDI-ToF dataset (a drug-dosed renal cancer model), we demonstrated SMEW’s generalisability and adaptability across diverse experimental designs, MSI technologies and biological contexts. Across these applications, SMEW enables users to perform structured comparisons between experimental conditions, identify spatially resolved metabolic patterns and assess how they evolve with disease or treatment, and prioritise regions of interest for detailed investigation, all within a code-free interactive interface.

Building on established MSI processing and statistical frameworks, SMEW provides a unified, hierarchical analysis workflow that integrates pseudobulk-, region-, and pixel-level analyses within a single environment. Statistical methods within SMEW are tailored to each analytical layer to accommodate substantial differences in sample size and to account for spatial dependence structures among neighbouring pixels. Importantly, the platform enables integration of multi-slide MSI datasets while preserving biologically meaningful spatial structures, allowing users to distinguish technical variation from condition-associated metabolic patterns. SMEW extends existing approaches by enabling spatially resolved pathway enrichment, metabolite co-localisation and covariation network analyses, as well as cross-sample and cross-condition spatial comparisons, and by supporting integration with transcriptomic and proteomic data. Apps created with SMEW are standalone and can be deployed online for sharing with collaborators and the community, bringing data directly to the hands of biologists without any installation or command line knowledge needed. The platform also supports export of pseudobulk intensity matrices and pixel-level metadata with newly derived clusters and annotations to enable further downstream analyses or integration with spatial multi-omic data, for example with MAGPIE^16^. Together, these capabilities position SMEW as a complementary framework facilitating advanced spatially informed metabolomics analysis, enabling robust biological interpretation of tissue heterogeneity and insights into disease progression and therapeutic response.

## Acknowledgements

We thank A. Borde and T. Volckaert for providing the mice from the bleomycin mouse model study, L. Setyo for providing histopathology annotations for the bleomycin mouse model, S. Hoffmann for providing the staining on liver ASO samples and J. Pfeiff for histological guidance on liver ASO samples.

## Data availability

SMEW is available on GitHub (https://github.com/Core-Bioinformatics/smew), with extensive documentation and examples at https://core-bioinformatics.github.io/smew. The bleomycin-treated mouse lung samples were previously published in Williams et al^16^, with raw data deposited in the MetaboLights database under accession code <u>MTBLS13445</u>. A live example app is also available at https://www.mohorianulab.org/shiny/smew/BLM_example/. The newly-generated mouse liver ASO data will be made available upon publication.

## Methods

### Pipeline Implementation and Usage

SMEW is implemented as an R package with a central function <u>create_smew_app()</u> which performs all required preprocessing, such as creating a pseudobulk intensity and metadata matrix, optional peak annotation and optional denoising. This function creates an R shiny app, with 4 overarching tabs, (1) an introductory tab which enables exploration of the peak annotation table and simple spatial visualisation, (2) a pseudobulk tab which features dataset-wide analysis at the whole sample level, (3) the region-level tab which allows users to identify regions of interest within each sample and (4) the pixel-level tab where colocalised sets of metabolite peaks and spatial areas showing high pathway activation can be studied. Further information on the individual panels is presented in the Results & Discussion section. The pipeline is flexible across MSI technologies, such as MALDI and DESI, and robust on the normalisation method used. SMEW can also incorporate histology images, which can be co-registered with MSI data using a provided Python app.

### Experimental methods

Experimental approaches for bleomycin-treated mice are described in Williams et al^16^.

For AZD2014/AZD8186 mice, experimental methods were described in Ling et al^6^.

For ASO-treated mice in the acute study, female BALB/c mice (6 weeks of age upon arrival) were acclimatised for 4 days then subjected to a single dose of SPC5001 (300 mg/kg), ASO1 (300 mg/kg) or saline. Liver lobes were collected on Day 3 post-dosing and snap-frozen in dry ice-cooled isopentane immediately after resection. Liver samples were then embedded in a hydrogel containing 7.5% hydroxypropyl methylcellulose (viscosity 40–60 cP, 2% in H2O at 20°C) and 2.5% polyvinylpyrrolidone as previously described^38^. Sections at 10 µm were cut on a CM1950 cryostat (Leica Biosystems, Nussloch, Germany), thaw-mounted onto Superfrost glass slides, dried under N2, vacuum-packed in slide mailers, and stored at −70°C. Slides were equilibrated to room temperature before unpacking for imaging. Desorption electrospray ionisation (DESI)-MSI was performed on an automated 2D DESI stage (Prosolia Inc., Indianapolis, USA) with a custom-built sprayer assembly^39^, coupled to a Q Exactive Plus (Thermo Scientific, Bremen, Germany). Data were acquired in negative ion mode (*m/z 80– 900*) and positive ion mode (*m/z 100–1000*) at a mass resolution of 70,000, with a fixed injection time of 150 ms and a spatial resolution of 55 µm. The electrospray solvent was MeOH/H2O (95:5, v/v) at 1.5 µL/min, with a spray voltage of ±4.5 kV and a nebulising gas pressure of 6 bar. Raw files were converted to .mzML using ProteoWizard msConvert^40^ (v.3.0.4043) and complied into imzML (imzML converter^41^ v1.3). Data were processed in SCiLS™ Lab MVS Premium 3D (v2026a, Bruker Daltonik, Bremen, Germany).

### Computational pre-processing

For all datasets, .imzml data files were initially imported into SCiLS Lab software (Bruker Daltonics, Germany, 2022a MVS). Bisecting k-means segmentation was performed within SCiLS Lab to create two regions of interest (ROIs): tissue and background (area outside the tissue). Individual samples were then segmented using the tissue/background ROIs with some manual alterations. Individual pixel-level total ion count (TIC) normalised peak intensities were extracted using the SCiLS Lab API, which yields a pixel-by-peak table with associated metadata. This table was then used as input to SMEW and peak annotation was performed through SMEW.

## Figures

**Supplementary Figure 1.**
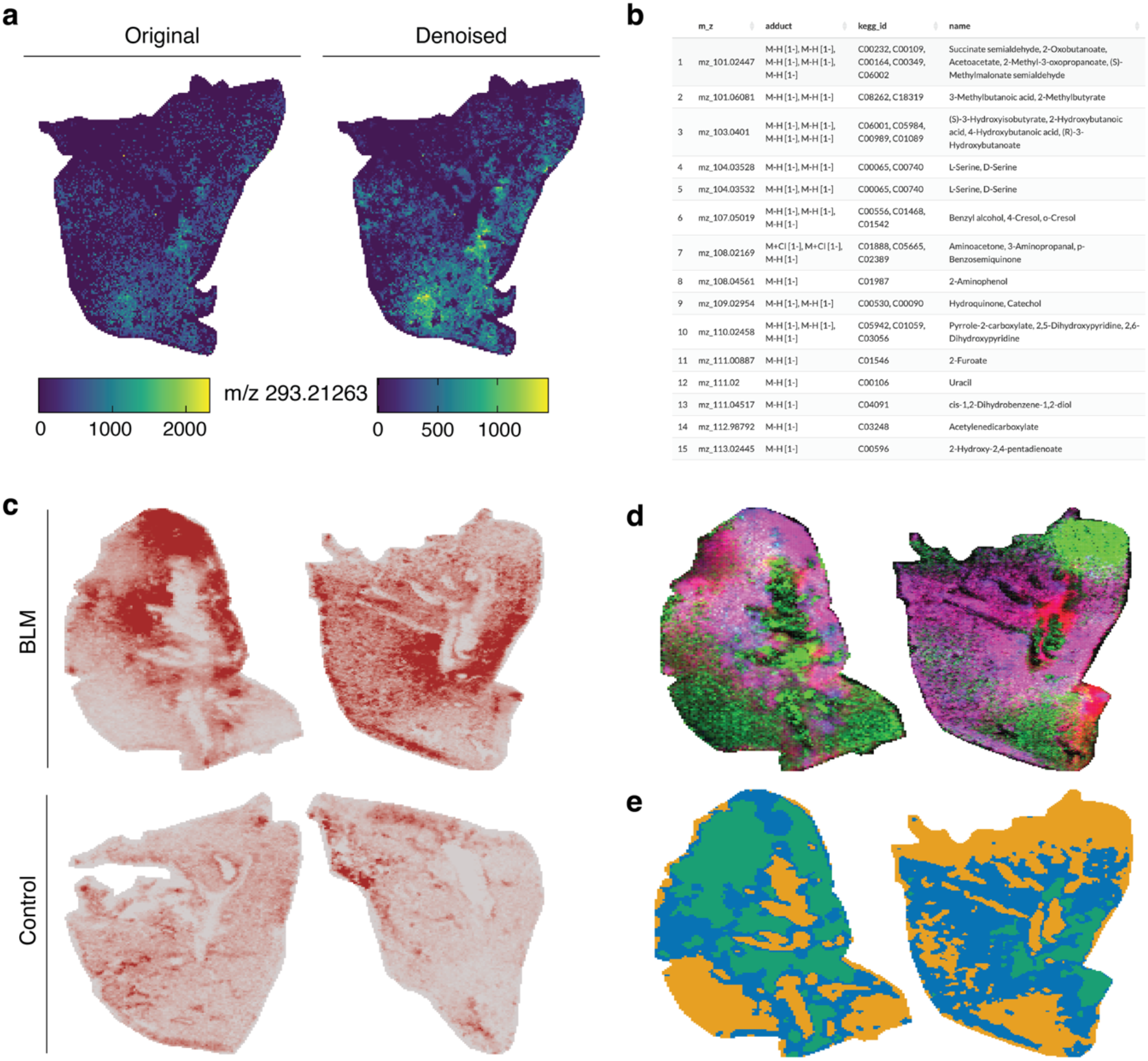
SMEW preprocessing and initial visualisation of bleomycin-induced lung fibrosis DESI-MSI dataset. (a) Example of spatial denoising applied to *m/z 293*.*21263*, showing the raw spatial intensity distribution (left) and the denoised image (right), illustrating improved signal coherence while preserving spatial structure. (b) Interactive annotation table within the SMEW interface, enabling users to explore peaks of interest. (c) Spatial visualisation of an individual metabolite peak (*m/z 303*.*23295*), across different tissue sections, demonstrating interactive inspection of spatial intensity patterns across the tissue section. (d) Composite RGB image showing the spatial variation of three selected metabolite peaks (*m/z 279*.*23196, m/z 89*.*02435, m/z 303*.*23295*), visualised as red, green, and blue channels to highlight co-localisation and regional heterogeneity. (e) Spatial distribution of user-provided pixel-level metadata (here, an external clustering), illustrating SMEW’s capacity for integration of external annotations (e.g. region labels or pathology classifications) for downstream spatial comparison and analysis.

**Supplementary Figure 2.**
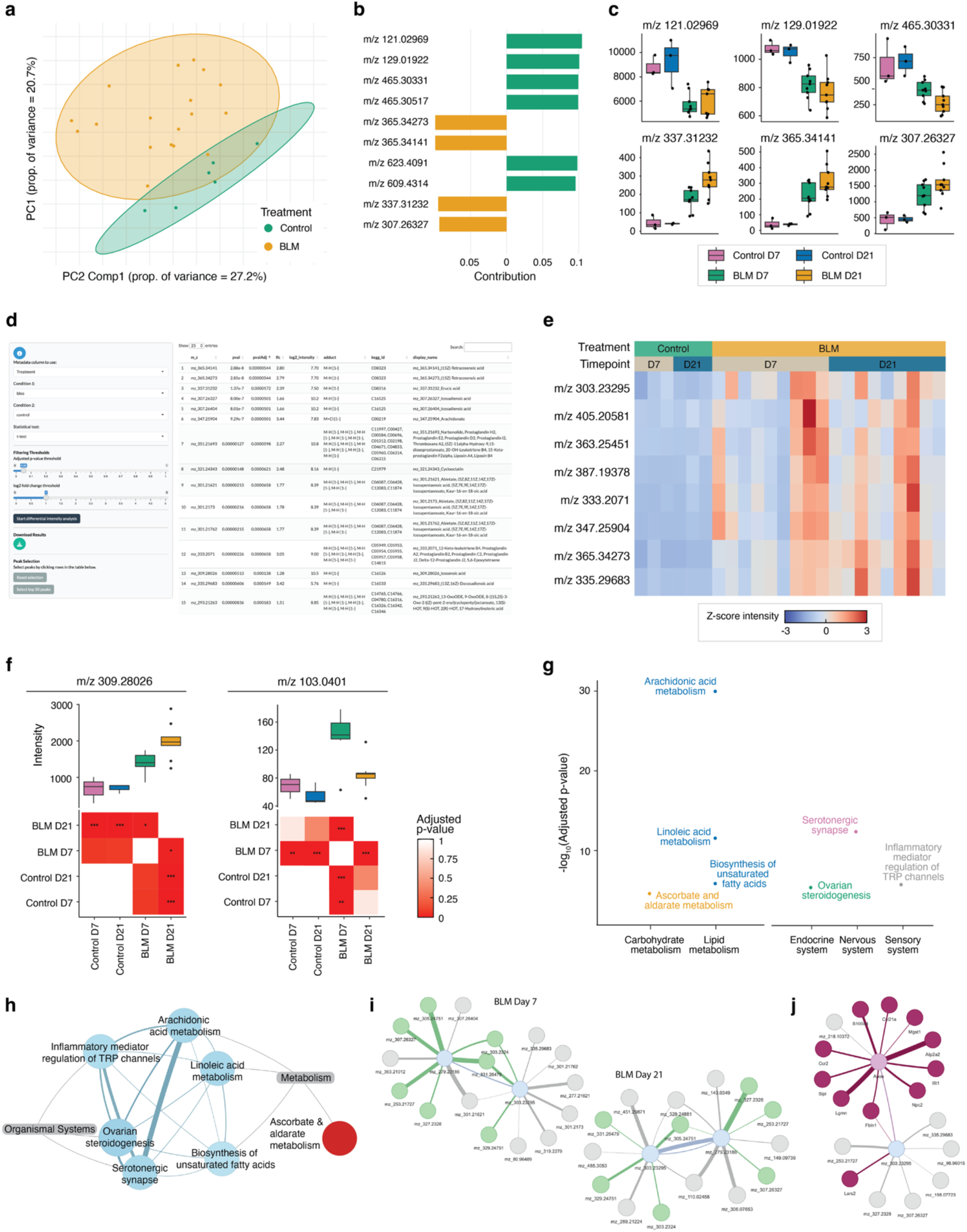
Bulk-level analysis workflow in SMEW applied to the bleomycin-induced lung fibrosis dataset. (a) Principal component analysis (PCA) of pseudobulk data, coloured by treatment group, illustrating global metabolic differences between control and bleomycin (BLM)-treated samples. (b) Top 10 feature contributions from partial least squares discriminant analysis (PLS-DA), using treatment group as the discriminative variable. Bars represent relative contribution of each metabolite peak to group separation. (a) Boxplots showing intensity distributions of selected top discriminative peaks across treatment and timepoint groups, highlighting condition-specific metabolic changes. (d) Screenshot of the differential intensity analysis module within the SMEW interface, enabling interactive selection of comparisons and statistical test. (e) Heatmap of Z-scaled intensities across samples for selected top differentially abundant peaks between control and BLM-treated groups, demonstrating coordinated metabolic shifts. Sample annotations are shown at the top. (f) Post-hoc Tukey’s test results following ANOVA comparison across treatment/timepoint groups for selected peaks. Upper panels show boxplots of peak intensities across groups; lower panels display pairwise Tukey’s test comparisons with Benjamini–Hochberg (BH) correction. Statistical significance is indicated as * p < 0.05, ** p < 0.01, and *** p < 0.001. (g) Enriched KEGG pathways derived from significantly altered peaks between control and BLM samples, visualised as −log_10_(BH adjusted *p*-value) and grouped by KEGG functional categories. (h) Pathway overlap network for significantly enriched KEGG pathways. Nodes represent pathways, edge weights indicate the number of shared metabolite peaks between pathways and nodes are additionally annotated by KEGG category. (i) GENIE3-based covariation networks constructed for BLM day 7 (left) and BLM day 21 (right) samples, using *m/z 303*.*23295* and *m/z 279*.*23186* as target features. Overlapping peaks between the two networks are highlighted in green, illustrating shared and timepoint-specific metabolic associations. (j) GENIE3-based covariation network on metabolomics and transcriptomics data from BLM day 21 samples using *Apoe* and *m/z 303*.*23295* as target features. Metabolite targets are shown in blue, gene targets in light pink, predicted metabolite regulators in grey and predicted gene regulators in purple.

**Supplementary Figure 3.**
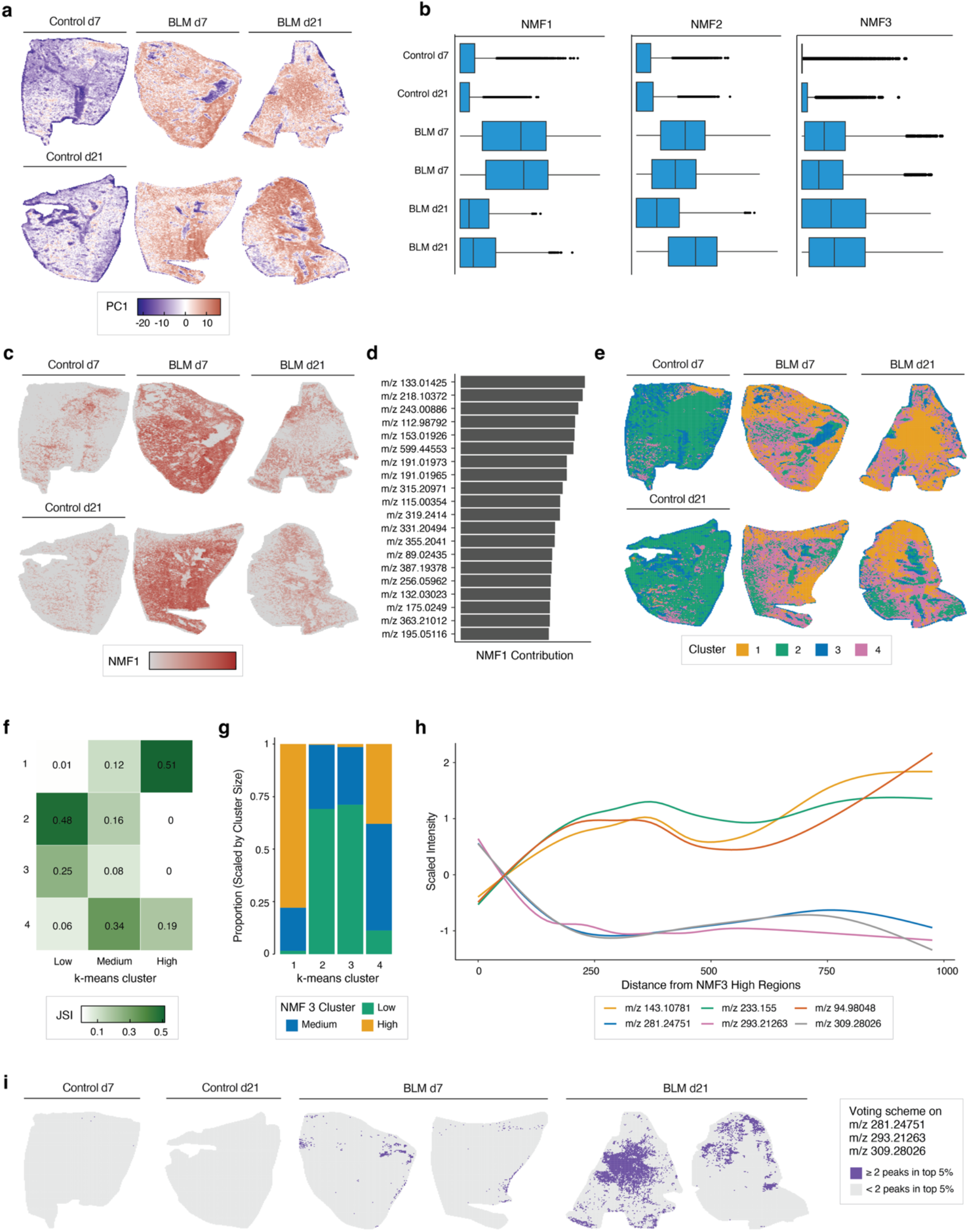
Region-level analysis workflow in SMEW applied to the bleomycin-induced lung fibrosis dataset. (a) Spatial visualisation of principal component 1 (PC1) across samples from different treatment groups, highlighting regional metabolic heterogeneity. (b) Boxplots showing variation in non-negative matrix factorisation (NMF) factors 1–3 across treatment and timepoint groups, illustrating condition-associated shifts in regional metabolic structure. (c) Spatial distribution of NMF1 across representative tissue sections, showing highest levels in day 7 BLM samples. (d) Bar plot of the top contributing metabolite peaks to NMF1, ranked by loading. (e) Spatial visualisation of k-means clustering (*k*=4) applied to pixel-level data, identifying metabolically distinct tissue regions. (f) Jaccard similarity index (JSI) comparison between NMF3-derived Low/Medium/High regions and k-means clusters, quantifying overlap of pixels between clustering approaches. (g) Bar plots showing the proportion of NMF3 regions (Low, Medium, High) within each k-means cluster, scaled by the size of each NMF3 region to illustrate relative enrichment. (h) Line plots depicting the variation in intensities of the top Pearson-correlated metabolite peaks as a function of distance from NMF3-high regions, demonstrating spatial gradients in associated features. (i) Spatial visualisation of a voting scheme based on the top negatively correlated peaks with increasing distance from NMF3-high regions, highlighting spatial domains specific to treatment groups, defined by coordinated metabolic variation.

**Supplementary Figure 4.**
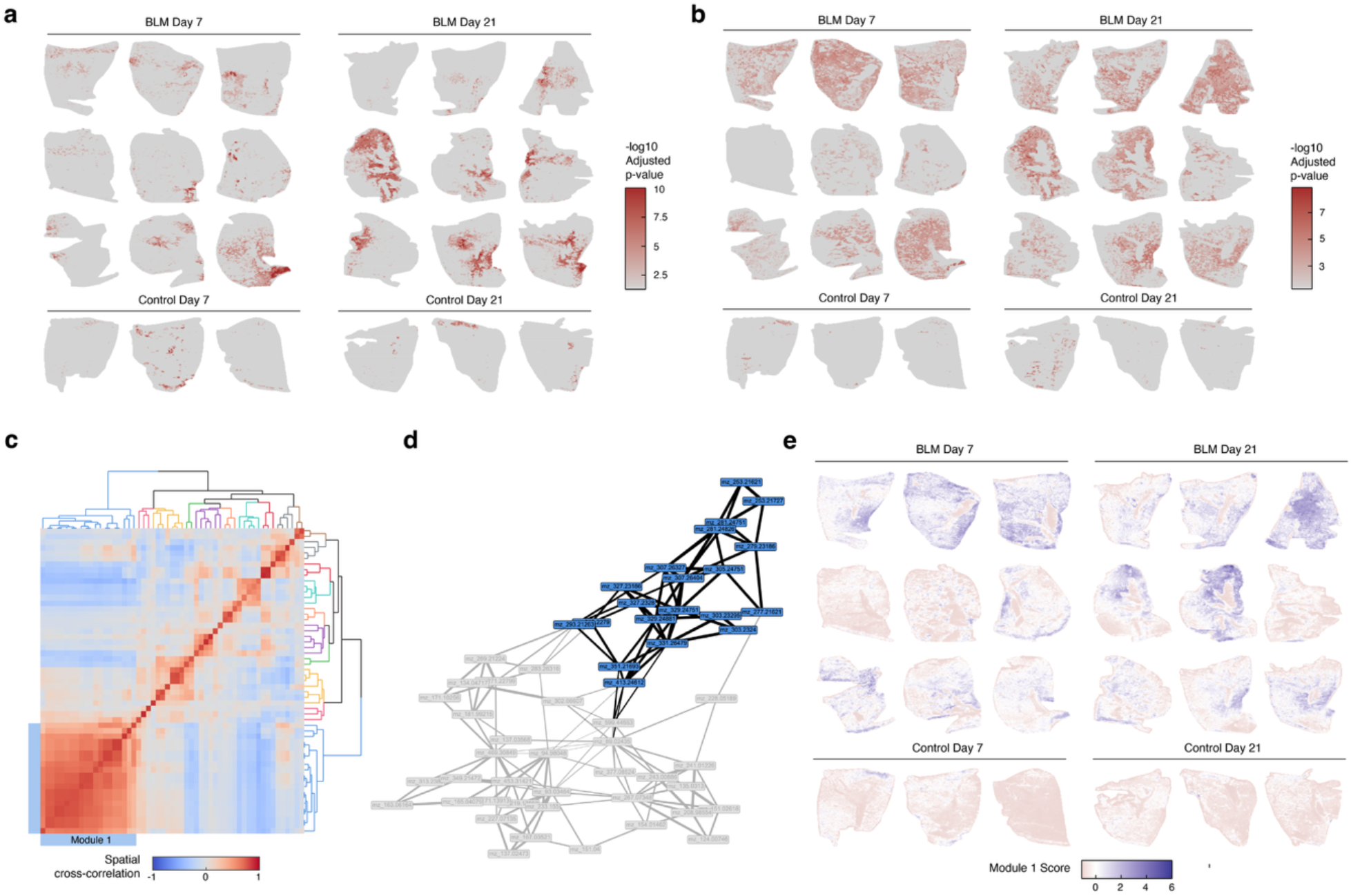
Pixel-level spatial analysis in SMEW applied to the bleomycin-induced lung fibrosis dataset. (a–b) Pixel-level spatial pathway enrichment analysis showing −log_10_(Benjamini Hochberg adjusted p-value) across the tissue section for (a) alpha-linoleic acid metabolism and (b) inflammatory mediator regulation of TRP channels pathways. Colour intensity reflects the local statistical significance of pathway enrichment derived from sliding-window–based neighbourhood comparisons. (c) Spatial cross-correlation heatmap of top autocorrelated (spatially variable) metabolite peaks, illustrating co-localisation patterns across pixels. Hierarchical clustering groups peaks into modules with shared spatial structure, with Module 1 highlighted. (d) Spatial cross-correlation network highlighting Module 1 (blue). Nodes represent metabolite peaks and edges denote top 3 spatial cross-correlations per peak. (e) Spatial visualisation of the Module 1 score across the tissue section, demonstrating coordinated regional enrichment of co-localised metabolite peaks specific to BLM samples.

**Supplementary Figure 5.**
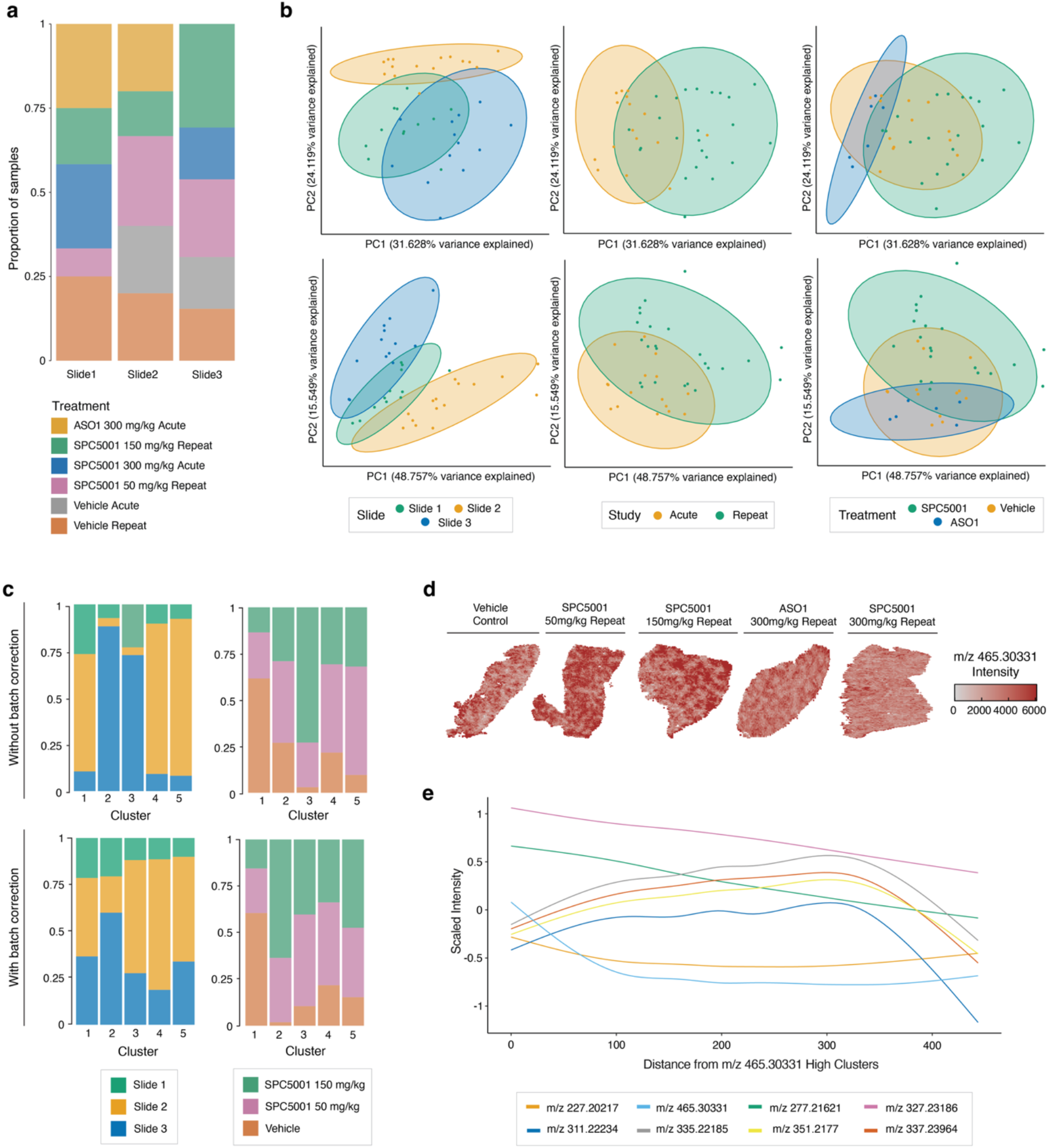
Bulk- and region-level analysis of the ASO-induced liver toxicity dataset using SMEW. (a) Distribution of samples across slides (batches) and treatment groups, illustrating the experimental batch structure of the dataset. (b) Bulk-level principal component analysis (PCA) of negative ion mode (top) and positive ion mode (bottom) data. Points are coloured by slide (left), study (middle), and treatment group (right), demonstrating sources of technical and biological variation. (c) Proportion of k-means clusters (*k*=5) assigned to each slide and treatment group without (top) and with (bottom) Harmony batch correction applied per slide, illustrating the effect of batch correction on clustering structure. (d) Distribution of *m/z 465*.*30331* intensity across treatment groups, highlighting zonation differences. (e) Radial distance analysis showing scaled metabolite intensities as a function of distance from *m/z 465*.*30331*-high regions, demonstrating spatial gradients associated with cholesterol sulphate-enriched periportal zones.

**Supplementary Figure 6.**
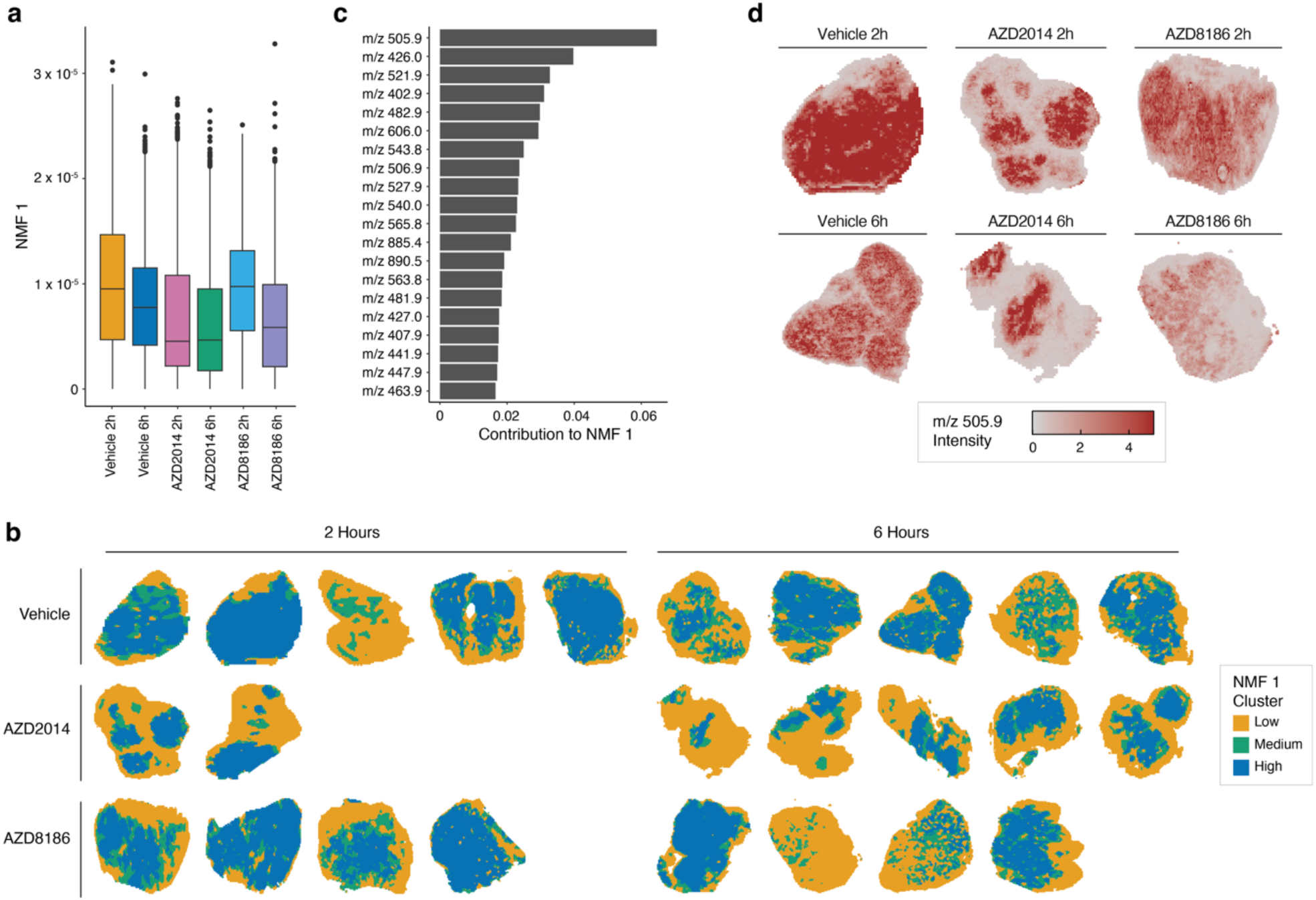
Analysis of small molecule-treated renal tumour MALDI-ToF dataset using SMEW. (a) Boxplots showing distribution of NMF Factor 1 scores per treatment group, highlighting treatment heterogeneity. (b) Spatial view of NMF Factor 1-based regions in selected samples, where “Low” corresponds to the bottom 40% and “High” the top 40% by NMF Factor 1. (c) Weights of top 20 metabolite peaks contributing to NMF Factor 1. (d) Spatial view of ATP (m/z 505.9) in selected samples.

## References

1 Körber, A., Anthony, I. G. M. & Heeren, R. M. A. Mass Spectrometry Imaging. Analytical Chemistry 97 (July 17, 2025). 10.1021/acs.analchem.4c05249

2 Buchberger, A. R., DeLaney, K., Johnson, J. & Li, L. Mass Spectrometry Imaging: A Review of Emerging Advancements and Future Insights. Analytical chemistry 90 (2017 Dec 13). 10.1021/acs.analchem.7b04733

3 B, F. et al. Investigation into Drug-Induced Liver Damage Using Multimodal Mass Spectrometry Imaging - PubMed. Journal of the American Society for Mass Spectrometry 36 (02/05/2025). 10.1021/jasms.4c00313

4 Sushentsev, N. et al. Spatial metabolomics informs the use of clinical imaging for improved detection of cribriform prostate cancer. Proceedings of the National Academy of Sciences 122 (2025-6-23). 10.1073/pnas.2502423122

5 Tsyben, A. et al. Cell-intrinsic metabolic phenotypes identified in patients with glioblastoma, using mass spectrometry imaging of 13C-labelled glucose metabolism. Nature Metabolism 7 (2025). 10.1038/s42255-025-01293-y

6 Ling, S. et al. Use of metabolic imaging to monitor heterogeneity of tumour response following therapeutic mTORC1/2 pathway inhibition. Disease Models & Mechanisms 18 (2025/02/01). 10.1242/dmm.050804

7 Samarah, L. Z. et al. Spatial metabolic gradients in the liver and small intestine. Nature 2025 648:8092 648 (2025-10-15). 10.1038/s41586-025-09616-5

8 Ràfols, P. et al. rMSI: an R package for MS imaging data handling and visualization. Bioinformatics 33 (2017/08/01). 10.1093/bioinformatics/btx182

9 Ràfols, P. et al. rMSIproc: an R package for mass spectrometry imaging data processing. Bioinformatics 36 (2020/06/11). 10.1093/bioinformatics/btaa142

10 Gibb, S. & Strimmer, K. MALDIquant: a versatile R package for the analysis of mass spectrometry data. Bioinformatics 28 (2012/09/01). 10.1093/bioinformatics/bts447

11 Saw, N. M. M. T. et al. MSI-Explorer: Napari-Powered Tool for Mass Spectrometry Imaging Data Analysis. Analytical Chemistry 97 (June 6, 2025). 10.1021/acs.analchem.5c01513

12 Föll, M. C. et al. Accessible and reproducible mass spectrometry imaging data analysis in Galaxy. GigaScience 8 (2019/12/01). 10.1093/gigascience/giz143

13 Mass Spectrometry Imaging Data Analysis with ShinyCardinal. (2024-03-12). 10.21203/rs.3.rs-4072606/v1

14 Moutsopoulos, I., Williams, E. C. & Mohorianu, I. I. bulkAnalyseR: an accessible, interactive pipeline for analysing and sharing bulk multi-modal sequencing data. Briefings in Bioinformatics 24 (2023/01/19). 10.1093/bib/bbac591

15 Franzen, L. et al. Mapping spatially resolved transcriptomes in human and mouse pulmonary fibrosis. Nat Genetics 56, 1725–1736 (2024). 10.1038/s41588-024-01819-2

16 Williams, E. C. et al. Spatially resolved integrative analysis of transcriptomic and metabolomic changes in tissue injury studies. Nature Communications 2026 17:1 17 (2026-01-07). 10.1038/s41467-025-68003-w

17 Katajamaa, M., Miettinen, J. & Orešič, M. MZmine: toolbox for processing and visualization of mass spectrometry based molecular profile data. Bioinformatics 22 (2006/03/01). 10.1093/bioinformatics/btk039

18 Bemis, K. A., Foll, M. C., Guo, D., Lakkimsetty, S. S. & Vitek, O. Cardinal v.3: a versatile open-source software for mass spectrometry imaging analysis. Nat Methods 20, 1883–1886 (2023). 10.1038/s41592-023-02070-z

19 Alexandrov, T. et al. Spatial Segmentation of Imaging Mass Spectrometry Data with Edge-Preserving Image Denoising and Clustering. (November 15, 2010). 10.1021/pr100734

20 Kanehisa, M., Furumichi, M., Sato, Y., Matsuura, Y. & Ishiguro-Watanabe, M. KEGG: biological systems database as a model of the real world. Nucleic Acids Research 53 (2025/01/06). 10.1093/nar/gkae909

21 Wishart, D. S. et al. HMDB 5.0: the Human Metabolome Database for 2022. Nucleic Acids Research 50 (2022/01/07). 10.1093/nar/gkab1062

22 Liebisch, G. et al. Update on LIPID MAPS classification, nomenclature, and shorthand notation for MS-derived lipid structures. Journal of Lipid Research 61 (2020/12/01). 10.1194/jlr.S120001025

23 Sementé, L., Baquer, G., García-Altares, M., Correig-Blanchar, X. & Ràfols, P. rMSIannotation: A peak annotation tool for mass spectrometry imaging based on the analysis of isotopic intensity ratios. Analytica Chimica Acta 1171 (2021/08/01). 10.1016/j.aca.2021.338669

24 Wadie, B. et al. METASPACE-ML: Context-specific metabolite annotation for imaging mass spectrometry using machine learning. Nature Communications 2024 15:1 15 (2024-10-22). 10.1038/s41467-024-52213-9

25 Weckerle, J. et al. Mapping the metabolomic and lipidomic changes in the bleomycin model of pulmonary fibrosis in young and aged mice. Dis Model Mech 15 (2022). 10.1242/dmm.049105

26 Oga, T. et al. Prostaglandin F2α receptor signaling facilitates bleomycin-induced pulmonary fibrosis independently of transforming growth factor-β. Nature Medicine 2009 15:12 15 (2009-11-29). 10.1038/nm.2066

27 Zhao, E. et al. Spatial transcriptomics at subspot resolution with BayesSpace. Nature Biotechnology 2021 39:11 39 (2021-06-03). 10.1038/s41587-021-00935-2

28 Korsunsky, I. et al. Fast, sensitive and accurate integration of single-cell data with Harmony. Nature Methods 2019 16:12 16 (2019-11-18). 10.1038/s41592-019-0619-0

29 Yuan, Z. et al. Benchmarking spatial clustering methods with spatially resolved transcriptomics data. Nature Methods 2024 21:4 21 (2024-03-15). 10.1038/s41592-024-02215-8

30 Shahsavari, A., Munteanu, A. & Mohorianu, I. ClustAssess: tools for assessing the robustness of single-cell clustering. bioRxiv (2022-02-02). 10.1101/2022.01.31.478592

31 Huynh-Thu, V. A., Irrthum, A., Wehenkel, L. & Geurts, P. Inferring regulatory networks from expression data using tree-based methods. PLoS One 5 (2010). 10.1371/journal.pone.0012776

32 Larsson, L., Franzen, L., Stahl, P. L. & Lundeberg, J. Semla: a versatile toolkit for spatially resolved transcriptomics analysis and visualization. Bioinformatics 39 (2023). 10.1093/bioinformatics/btad626

33 Miller, B. F., Bambah-Mukku, D., Dulac, C., Zhuang, X. & Fan, J. Characterizing spatial gene expression heterogeneity in spatially resolved single-cell transcriptomic data with nonuniform cellular densities. Genome Res 31, 1843–1855 (2021). 10.1101/gr.271288.120

34 Andersson, P. & den Besten, C. Preclinical and Clinical Drug-metabolism, Pharmacokinetics and Safety of Therapeutic Oligonucleotides. Advances in Nucleic Acid Therapeutics (2019/2/08). 10.1039/9781788015714-00474

35 A, G. et al. Considerations in the Preclinical Assessment of the Safety of Antisense Oligonucleotides - PubMed. Nucleic acid therapeutics 33 (2023 Jan). 10.1089/nat.2022.0061

36 Poelgeest, E. P. v. et al. Acute Kidney Injury During Therapy With an Antisense Oligonucleotide Directed Against PCSK9. American Journal of Kidney Diseases 62 (2013/10/01). 10.1053/j.ajkd.2013.02.359

37 Hall, Z. et al. Lipid zonation and phospholipid remodeling in nonalcoholic fatty liver disease. Hepatology 65 (April 2017). 10.1002/hep.28953

38 Dannhorn, A. et al. Universal Sample Preparation Unlocking Multimodal Molecular Tissue Imaging. Analytical Chemistry 92 (June 10, 2020). 10.1021/acs.analchem.0c00826

39 Takáts, Z. n., Wiseman, J. M., Gologan, B. & Cooks, R. G. Mass Spectrometry Sampling Under Ambient Conditions with Desorption Electrospray Ionization. Science 306 (2004-10-15). 10.1126/science.1104404

40 Adusumilli, R. & Mallick, P. Data Conversion with ProteoWizard msConvert. Methods in Molecular Biology (2017). 10.1007/978-1-4939-6747-6_23

41 Race, A. M., Styles, I. B. & Bunch, J. Inclusive sharing of mass spectrometry imaging data requires a converter for all. Journal of Proteomics 75 (2012/08/30). 10.1016/j.jprot.2012.05.035

